# Integrative k-mer and Transcriptomic Analysis Reveals Putative Sex-Determining Genes in *Spinacia turkestanica*

**DOI:** 10.1101/2025.10.22.683888

**Authors:** Shoaib Muhammad, Shuyu Liang, Tongyu Zhou, Xinrong Liu, Liang Yang, Ameer Ahmed Mirbahar, Ning Li, Chuanliang Deng

**Author notes:** Author for Correspondence: Chuanliang Deng,; Ning Li.

## Abstract

Deciphering the regulation of sex-determining gene/s in dioecious crops is crucial for molecular breeding. However, the identification of sex-determining gene/s in *Spinacia* is challenging due to incomplete genome assemblies, high genomic similarity between males and females, and the limitations of transcriptome-only analyses, which may miss unannotated or novel genes in poorly assembled or absent genomic regions. To overcome these limitations, we employed a reference-genome-free k-mer approach to identify candidate sex-determining genes in *Spinacia turkestanica*, the closest evolutionary relative of cultivated spinach (*S. oleracea*). Male-specific reads were *de novo* assembled into contigs, revealing 21.5 Mb of the sex-determining region (SDR). Using the MAKER pipeline, which integrates transcriptomic and proteomic evidence, we predicted 226 protein-coding genes within the SDR, including nine previously unannotated. Transcriptomic profiling combined with weighted gene co-expression network analysis identified eight SDR DEGs, including two newly annotated genes, co-expressing during early male flower development. qPCR validation confirmed three SDR genes as candidate sex-determining factors, including *TU_SDR00087* (bZIP domain), *TU_SDR000168* (RNA-binding splicing factor domain), and *TU_SDR000174* (MYB domain). Together, these findings provide a foundation for functional characterization of sex determination in *Spinacia*.

**Highlight:** A k-mer based analysis uncovered male-specific regions and candidate sex-determining genes in *Spinacia turkestanica*, advancing understanding of sex regulation and spinach breeding.

## Introduction

Dioecy, a reproductive system in which plant species have distinct male and female individuals, occurs in approximately 6% of angiosperms and represents a notable divergence from the more common hermaphroditic reproduction (Renner, 2014). In dioecious plants, two primary theories explain sex determination: the two-gene mutation model and the single-gene mutation model. The former involves the interaction of an *M* gene (controlling stamen fertility) and an *F* gene (regulating pistil development), while the latter proposes that mutations in a *Q* gene can independently induce male or female sterility (Henry *et al*., 2018; Barrett, 2002; Charlesworth, 2015; Feng *et al*., 2020).

Many dioecious plants, including date palm , papaya, yam, spinach, kiwifruit, hemp, ginkgo, and poplar hold agronomic, medicinal, and industrial significance (Torres *et al*., 2018; Liao *et al*., 2020; Xue *et al*., 2020; Akagi *et al*., 2023; She *et al*., 2023; Qiao *et al*., 2024; Shi *et al*., 2025; Chae *et al*.). Substantial progress has been made in identifying Sex Determining Regions (SDRs) and candidate sex-determining genes in these species. For example, in persimmon (*Diospyros lotus*), the Y-encoded *OGI* pseudogene suppresses the autosomal transcription factor *MeGI*, promoting male flower development (Akagi *et al*., 2014). Similarly, in date palm, a male-specific region harboring *CYP703* and *GPAT3* suggests their involvement in male floral organ formation (Torres *et al*., 2018). A parallel mechanism occurs in kiwifruit, where the Y-linked *FrBy* (Friendly Boy) gene activates stamen development, while *SyGl* (Shy Girl) independently suppresses female traits (Akagi *et al*., 2019). Recent genome assembly of asparagus (*Asparagus officinalis*) identified a putative male-promoting factor (*aspTDF1*) and the SUPPRESSOR OF FEMALE FUNCTION (*SOFF*) gene, which independently inhibit gynoecium and promote androecium development (Harkess *et al*., 2020). In poplar, male floral development is triggered by the suppression of *ARR17* orthologs by an SDR-encoded duplicate (Müller *et al*., 2020). Meanwhile, hemp studies implicate transcription factors from the REM, bZIP, and MADS-box families in sex determination (Shi *et al*., 2025).

Spinach (*Spinacia oleracea*) is the most widely cultivated leafy vegetable in China (Han *et al*., 2023*b*). The genus *Spinacia* comprises three species: the cultivated

*S. oleracea* and its two wild relatives, *S. turkestanica* and *S. tetrandra* (She *et al*., 2022). Comparative genomic analyses suggest that *S. turkestanica* is the direct progenitor of cultivated spinach, making it an ideal system to study both the genetic basis of sex determination and the evolution of sexual dimorphism during spinach domestication (She *et al*., 2025).

Multiple approaches have been employed to localize the Sex Determining Region (SDR) in *S. oleracea*. Yu et al. (2021) mapped the SDR to 45.2 cM on Linkage Group 1 (LG1) in the cultivar ’Cornell-NO. 9’ using resequencing-based genotyping of an F₁ population. In contrast, Ma et al. (2022) identified a 17.42 Mb SDR on chromosome 1 through integrated genetic mapping and genome-wide association studies, while She et al., (2023) proposed a distinct 24.1 Mb SDR on chromosome 4. Recently, a draft male *S. turkestanica* genome was assembled using *S. oleracea* as a reference, though many contigs remained unanchored to chromosomes (She *et al*., 2025). Our lab further resolved the SDR by sequencing the X and Y chromosomes of *S. turkestanica* via single-chromosome sequencing technology (Li *et al*., 2023). However, this assembly also relied on the *S. oleracea* reference genome, limiting its contiguity.

Comparative transcriptomic analyses have previously been used to predict candidate genes involved in floral organ development and sexual differentiation in *S. oleracea* (Li *et al*., 2022; Ma *et al*., 2022; You *et al*., 2025). However, transcriptome-based approaches alone may overlook unannotated or novel genes located in incompletely assembled regions or absent from reference genomes (Zhang *et al*., 2020). These limitations may partly explain why the master regulator of sex determination in genus *Spinacia* remains unidentified.

To overcome these limitations, we applied a reference-independent k-mer approach to capture male-specific reads in *S. turkestanica* and assembled them into SDR-associated contigs. Structural annotation revealed nine novel gene models within the SDR. By integrating these results with transcriptomic profiling and weighted co-expression analysis, we identified three strong candidate sex-determining genes: *TU_SDR00087* (bZIP domain), *TU_SDR000174* (MYB domain), and *TU_SDR000168* (RNA-binding splicing factor).

## Materials and Methods

### Genomic libraries and k-mer analysis

Genomic libraries for *S. turkestanica* (male = 19, female = 12) were obtained from the Institute of Vegetables and Flowers (IVF), Chinese Academy of Agricultural Sciences (CAAS) (She et al., 2024). Quality-trimmed paired-end reads (PE-reads) (Chen *et al*., 2018) were pooled by sex and processed into 31-bp k-mers using Jellyfish (Marçais and Kingsford, 2011), retaining those with a minimum count of 10. Male-specific k-mers (MSKs) were defined as absent in females, and female-specific k-mers (FSKs) as absent in males (Liao *et al*., 2020). PE-reads containing at-least one sex-specific k-mer were termed as sex-specific PE-reads.

Male-specific PE reads, extracted in canonical form using BBTools (Bushnell *et al*., 2017), were mapped to *S. turkestanica* Tu17S31XY (She *et al*., 2025) genome using BWA-mem2 (Vasimuddin *et al*., 2019) to identify candidate regions likely representing the sex-determining region (SDR). PE-reads mapped to these regions were then subjected to *de novo* assembly. Mapping depth was calculated in 1-Mb sliding windows using SAMtools (Li *et al*., 2009) after deduplication.

### *De novo* contig assembly

We generated high-quality contig assembly using SOAPdenovo2 (SOAPdenovo-127mer all -K 61 -m 127 -R -E -F -V -M 2 -D 2) (Luo *et al*., 2012) and SPAdes (-k 51,61,71,81,91,101,111,121 --isolate) (Prjibelski *et al*., 2020), leveraging their complementary strengths in resolving complex genomic regions and coverage variability. The resulting assemblies were merged and gap-closed using Redundans (Pryszcz and Gabaldón, 2016) pipeline. The remaining gaps were filled using Sealer (- b 40G -k90 -k80 -k70 -k60 -k50 -k40 -k30 -L 199 -G 3000) (Paulino *et al*., 2015), followed by three rounds of genome polishing with Pilon (Walker *et al*., 2014). The male-specific PE-reads were mapped to the newly assembled contigs to evaluate their male specificity.

### GWAS analysis

Paired-end genomic libraries were aligned to the *S. turkestanica* (Tu17S31XY) genome (She *et al*., 2025) using BWA-mem2 (Vasimuddin *et al*., 2019) with default parameters. Mapped reads were sorted and deduplicated with SAMtools (Li *et al*., 2009). Variants were called using BCFtools (Danecek *et al*., 2021) and filtered for quality scores ≥30, minor allele frequency (MAF) ≥0.05, and genotype rates ≥0.3. Genome-wide association studies (GWAS) were performed using GEMMA (Zhou and Stephens, 2012) to identify sex-associated SNPs. Significant SNP regions were visualized as Manhattan plots generated using a custom Python script.

### Gene model prediction and annotation

Repetitive elements in the assembled SDR contigs were masked using RepeatMasker (http://www.repeatmasker.org/RepeatMasker) with RepBase28.07 (Bao *et al*., 2015), Dfam (Storer *et al*., 2021), and publicly available spinach-specific repeat libraries (Xu *et al*., 2017; Li *et al*., 2021). Gene prediction was carried out using three iterative rounds of the self-trained MAKER (Cantarel *et al*., 2008) pipeline, integrating *ab initio*, transcriptomic, and proteomic evidence. For *ab initio* prediction, SNAP (Korf, 2004) and AUGUSTUS (Stanke *et al*., 2006) were used. Transcriptomic evidence was derived from RNA-seq data generated in this study and from publicly available datasets (PRJNA630139, PRJNA649901, PRJNA663445, PRJNA716151, PRJNA1062132, PRJNA724923), which were aligned to the SDR contigs using Hisat2 (Kim *et al*., 2019) and StringTie (Pertea *et al*., 2015). For proteomic evidence, protein sequences from *Arabidopsis*, *Beta vulgaris*, *Chenopodium quinoa*, multiple *S. oleracea* accessions, *S. tetrandra*, *S. turkestanica*, and Swiss-Prot were aligned to SDR contigs using Exonerate (https://github.com/nathanweeks/exonerate).

Predicted gene models were retained as high-confidence if they satisfied the following criteria: intact open reading frames (ORFs ≥ 40 amino acids), presence of conserved protein domains and transcriptomic support (TPM ≥ 1). Functional annotation was performed using NCBI CDD, InterProScan, UniProt, and TAIR database (Rhee *et al*., 2003; Marchler-Bauer *et al*., 2015; Cantalapiedra *et al*., 2021; Blum *et al*., 2025), and human-readable descriptions were assigned using the AHRD pipeline (https://github.com/groupschoof/AHRD).

Orthologs were identified through reciprocal BLASTp searches (E-value < 1e−5) using TBtools v2.0 (Chen *et al*., 2020) against the Tu17S31XY genome. To detect potentially unannotated genes, additional BLASTn searches (E-value < 1e−10) were performed against the Tu17S31XY genome using the newly predicted gene models.

### Transcriptome sequencing and RNA-seq analysis

Male and female flower samples of *Spinacia turkestanica*, cultivated under controlled conditions, were collected at five distinct developmental stages (S1–S5), as defined by Ma et al. (2022). Each stage was characterized based on floral bud size and validated using scanning electron microscopy (SEM) prior to sampling: Stage 1, buds 0.2–0.5 mm; Stage 2, buds 0.5–1 mm; Stage 3, emergence of sepal primordia and stamen initiation in male flowers, and ovarian dome differentiation in female flowers; Stage 4, anther development in males and pistil formation with ovule initiation in females; Stage 5, anther maturation in males and ovule maturation with stigma emergence in females (Figure S5)**Error! Reference source not found.**. A total of 30 RNA-seq libraries (2 sexes × 5 stages × 3 biological replicates) were sequenced on the DNBSEQ-T7 platform, generating 150-bp paired-end reads (Table S8). Quality filtering was conducted using Fastp (Chen *et al*., 2018), and potential contamination from non-target species was removed using Kraken2 (Wood *et al*., 2019). Ribosomal RNA was filtered out using SortMeRNA (Kopylova *et al*., 2012) with the Rfam (Ontiveros-Palacios *et al*., 2025) and SILVA (Quast *et al*., 2013) databases to enrich for coding sequences. Cleaned reads were mapped to the *de novo* assembled contigs using HISAT2 (Kim *et al*., 2019), and gene expression levels were quantified with FeatureCounts (Liao *et al*., 2014). Differential expression analysis was performed using DESeq2 (Love *et al*., 2014) with default settings. Genes exhibiting a ≥2-fold change and an adjusted p-value ≤ 0.01 between male and female flowers at any stage were considered differentially expressed genes (DEGs).

### Weighted gene co-expression network analysis (WGCNA)

Genes with TPM > 1 in at least one sample were log2-transformed [log2(TPM + 1)] for variance stabilization, and the median absolute deviation (MAD) was calculated for each gene across samples. Based on MAD scores, the top 50% of genes, including DEGs (total n = 9,842), were used to construct a weighted gene co-expression network using PyWGCNA (Rezaie *et al*., 2023). To achieve a scale-free topology (R² > 0.85), a soft-threshold power of 15 was applied for a signed network, with a minimum module size of 30 and a deepsplit of 3. Module–trait correlations were computed to identify gene clusters associated with early-stage flower development. Genes in the target modules were functionally annotated using InterProScan, UniProt, and TAIR databases. Hub genes were selected based on kME (> 0.8) and intramodular connectivity (kWithin), and edges with weights > 0.2 were visualized in Cytoscape (Shannon *et al*., 2003) to depict hub gene network connectivity.

### Functional enrichment analysis

GO terms for the *S. turkestanica* genome were assigned using a Blast2GO-style pipeline that integrated homology and domain-based annotations from eggNOG-mapper, InterProScan, UniProt, and TAIR (Rhee *et al*., 2003; Cantalapiedra *et al*., 2021; Blum *et al*., 2025; The UniProt Consortium, 2025). This approach ensured broad and accurate functional coverage across the gene set. The consolidated annotations were then used for enrichment analysis of target genes with TBtools (Chen *et al*., 2020).

### qPCR analysis

To validate the differentially expressed genes (DEGs) identified between male and female flowers, qPCR analysis was performed using the same RNA samples utilized for RNA-seq library preparation. First-strand cDNA was synthesized with the PrimeScript™ RT reagent Kit with gDNA Eraser (TaKaRa) to eliminate genomic DNA contamination. Each qPCR reaction was prepared using TB Green® Premix Ex Taq™ II (Tli RNaseH Plus) (TaKaRa), with *spUBI* serving as the internal reference gene. Reactions were run on the LightCycler® 480 System (Roche) following the manufacturer’s protocol. Three biological replicates and two technical replicates were included to ensure reproducibility and minimize experimental error. Relative gene expression levels were calculated using the 2^−ΔCt method, and standard deviation (SD) was derived from the biological replicates. The primer sequences used for qPCR are listed in Table S17.

### SEM observation

Scanning electron microscopy (SEM) was employed to standardize the five developmental stages of male and female flowers in *Spinacia turkestanica*, utilizing a previously established protocol in our laboratory (Li *et al*., 2025). Floral samples were collected at distinct developmental stages and fixed in FAA solution (50% ethanol, 5% glacial acetic acid, and 1.85% formaldehyde; v/v/v), followed by vacuum infiltration for 10 minutes. After 4 hours, the fixative was replaced, and samples were incubated overnight at 4 °C. Dehydration was carried out using an ethanol gradient (50%, 70%, 80%, 90%, and three changes of 100%), with each step lasting 25 minutes. A final dehydration step was performed in tert-butanol for 20 minutes. The samples were then rapidly frozen in liquid nitrogen for 5 minutes and freeze-dried under vacuum for 2–3 hours. Gold sputter coating was applied at 10 mA for 20 seconds per cycle, with 1-minute intervals between cycles, repeated 2–3 times. SEM imaging was conducted using a Hitachi TM3030 Plus microscope to document and define the floral developmental stages.

## Results

The published *S. turkestanica* genome represents an XY male, which complicates the identification of the male-specific sex-determining region (SDR) through comparative genomics due to the lack of an XX female reference. To overcome this limitation, we developed a k-mer–based pipeline (**Figure 6**) to extract male-specific reads, assemble them *de novo* into SDR sequences, and predict candidate sex-determining genes.

**Figure 1.** Schematic of k-mer based approach to detect male-specific contigs and DEGs candidate for sex determination in *S. turkestanica*.

**Figure 2.** Identification and validation of *S. turkestanica* sex determining region.

**Figure 3.** Temporal expression of *de novo* assembled SDR gene models in reproductive tissues.

**Figure 4.** Gene expression dynamics of *S. turkestanica* male and female flowers.

**Figure 5.** Weighted co-expression network analysis of *S. turkestanica*.

**Figure 6.** Expression analysis of selected SDR genes across five developmental stages in *S. turkestanica* male and female flowers.

**Figure 6.**
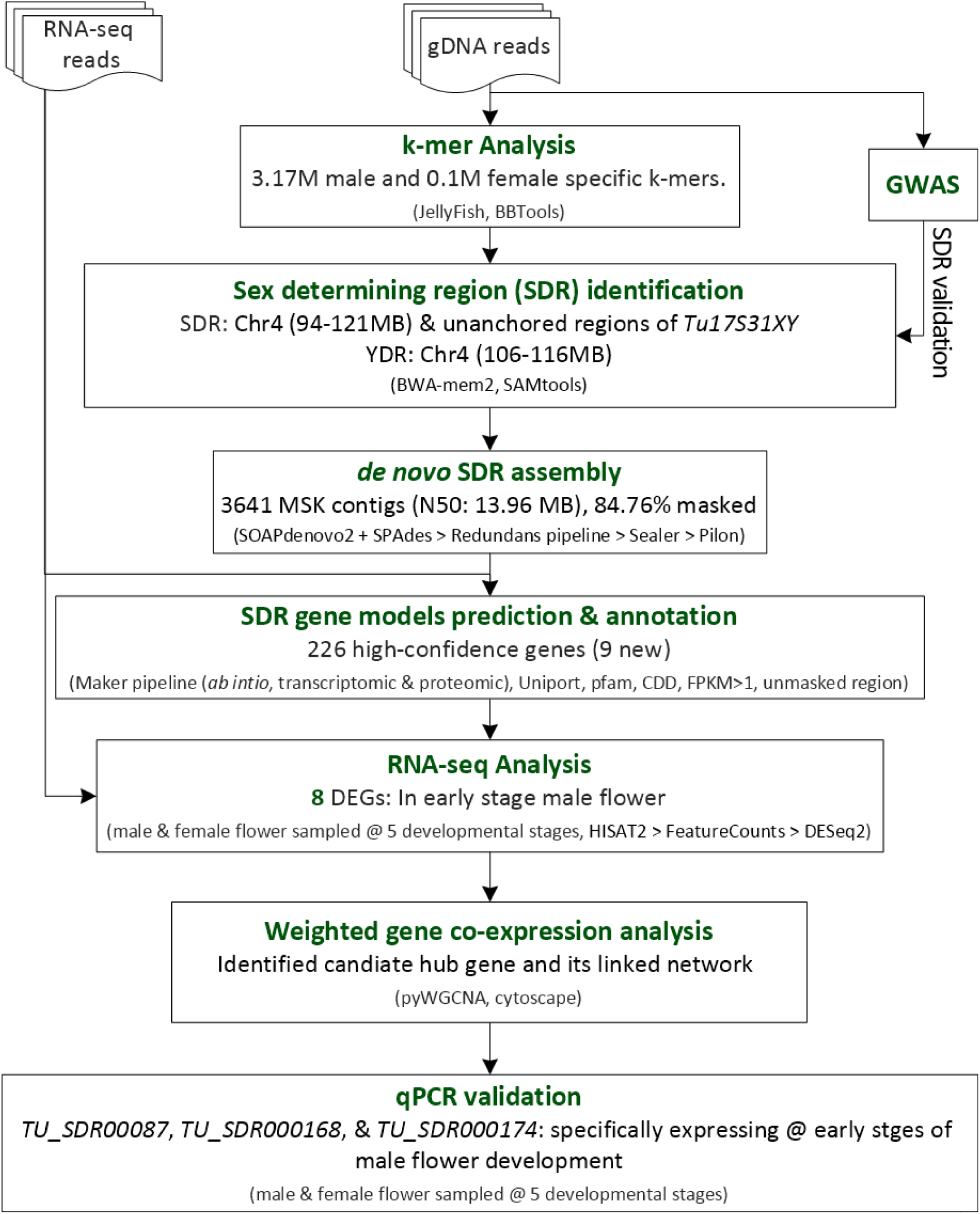
Schematic of k-mer based approach to detect male-specific contigs and DEGs candidate for sex determination in *S. turkestanica*.

### XY sex determination system in *S. turkestanica* centers on chromosome 4

We cataloged the paired-end (PE) reads, from the *S. turkestanica* male (n=19) and female (n=12) genomic groups, into 31 bp k-mers and identified 3,179,358 male-specific (MSKs) and 107,627 female-specific k-mers (FSKs) after PE-read normalization. The number of MSKs was approximately 29-fold higher than that of FSKs, indicating the existence of male-specific genomic regions in *S. turkestanica* (Figure 7a).

**Figure 7.**
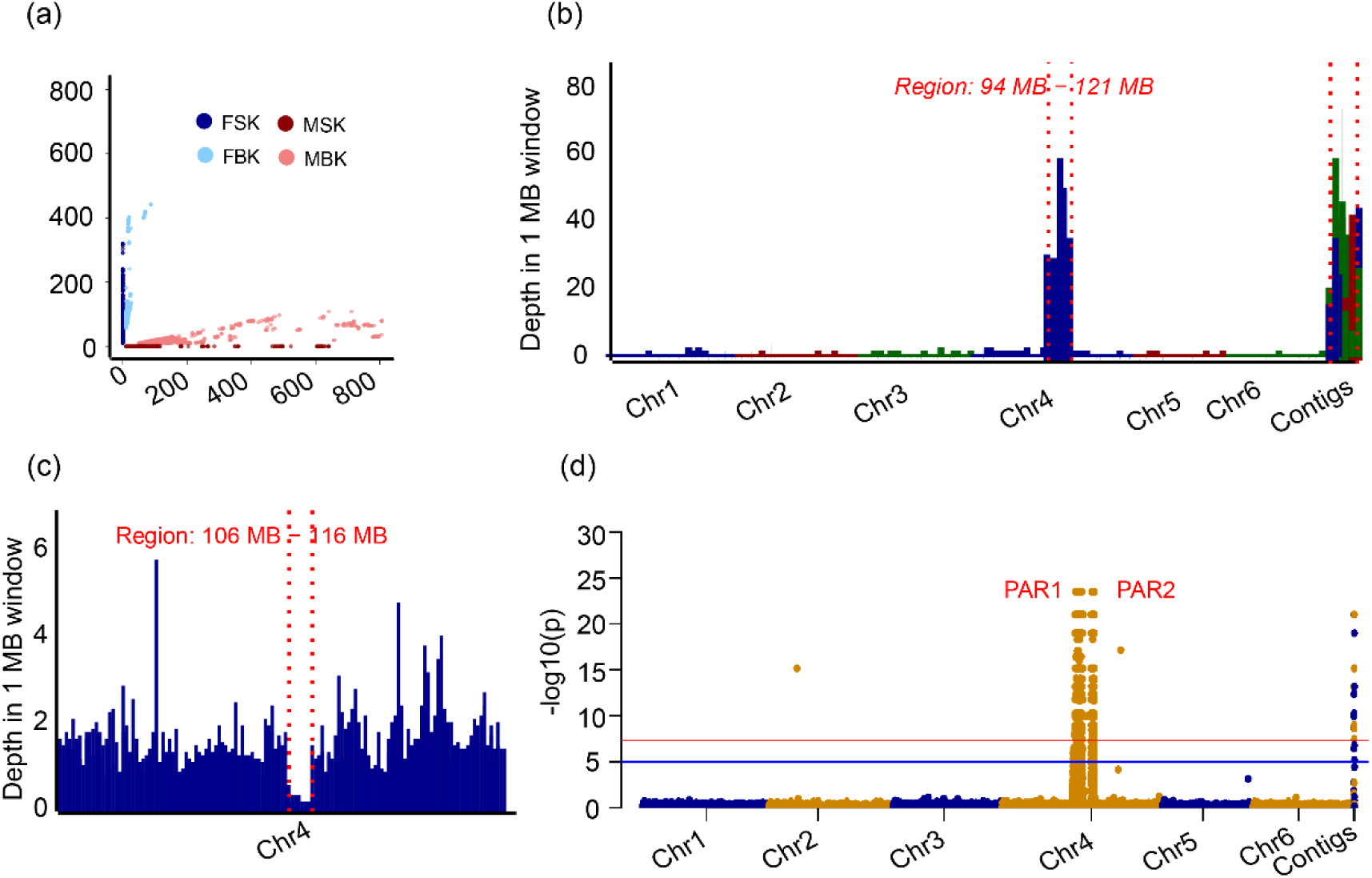
Identification and validation of *S. turkestanica* sex determining region. (a) Distribution of sex-specific k-mers in male and female genome. FSK, female-specific k-mer; MSK, male-specific k-mer; FBK, female-biased k-mer; MBK, male-biased k-mer. (b) Mapping of male-specific reads to the *Tu17S31XY* published genome. Vertical red dotted lines mark the core sex-determining regions on Chromosome 4 and unanchored contigs (c) Mapping of female-specific reads to the *Tu17S31XY* Chr 4. Vertical red dotted lines mark the core Y-duplication regions on Chromosome 4. Each bar represents the average depth in the 1-Mb sliding window. (d) Manhattan plot based on the genome-wide association studies (GWAS) using 19 male and 12 female accessions. Peaks on chromosome 4 indicate the proposed pseudo-autosomal regions. The red horizontal line indicates the significant threshold at α < 0.05.

In *S. oleracea*, the male-specific genomic region, or sex-determining region (SDR), is subdivided into the Y-duplication region (YDR) and two pseudo-autosomal regions (PAR1 and PAR2) (She *et al*., 2023). To determine whether *S. turkestanica* exhibits a similar SDR organization, we mapped 15,169,111 male-specific and 7,051,051 female-specific paired-end (PE) reads to the Tu17S31XY male genome (She *et al*., 2025). The majority of male-specific reads (95.35%) mapped to the reference, with 40.39% concentrated in the 94–121 Mb interval of chromosome 4, indicating this region as the SDR (Figure 7b). An additional 16.11% aligned to unplaced contigs (Table S1), likely representing SDR sequences currently unanchored in published genome. In contrast, female-specific reads were evenly distributed across the genome, except for a 10 Mb region (106–116 Mb) on chromosome 4 showing no coverage, suggesting a Y-duplication region (YDR) within the *S. turkestanica* SDR (Figure 7c, Figure S1a).

To validate our k-mer–based findings, we performed a genome-wide association study (GWAS), identifying 1,957 SNPs significantly associated with sex. Of these, 1,937 SNPs mapped to the 94–121 Mb interval on chromosome 4, forming two peaks (PAR1 and PAR2) flanking the proposed YDR, while 18 SNPs mapped to unanchored contigs (Figure 7d, Table S2). Nearly all SNPs were heterozygous in males (99.2%) and homozygous in females (99.7%) (Figure S1b). Together, these results indicate that the SDR in *S. turkestanica* resides on chromosome 4 and are consistent with an XY mechanism of sex determination.

### High-confidence gene set identified in the 21.5 Mb SDR of *S. turkestanica*

To further characterize the sex-determining region (SDR), we performed *de novo* assembly of male-specific paired-end reads, generating 3,641 male-specific contigs (N50: 13.96 MB) (Table S3), with the longest contig measuring 13.96 Mb and the shortest 200 bp. These contigs collectively spanned 21.5 Mb, of which 84.76% comprised repetitive elements, including 18.71% LTR/Copia and 24.06% LTR/Gypsy sequences (Table S4).

Using the MAKER annotation pipeline (Cantarel *et al*., 2008), we predicted 226 protein-coding gene models on the male-specific contigs (Table S5), with an average of 4.2 exons per transcript and a gene density of 8.5 genes per Mb (Table S6). While 305 genes have been annotated in the *S. turkestanica* (Tu17S31XY) SDR (She *et al*., 2025), our stringent annotation, filtered for RNA-seq support, protein domain conservation, and exclusion of repetitive regions, retained only 226 high-confidence genes (Table S5).

### New gene models identified and localized within the SDR structure

To assess conservation, we performed reciprocal BLASTp and supporting BLASTn searches against the *S. turkestanica* Tu17S31XY genome, identifying nine previously unannotated gene models within the SDR. Of 226 predicted genes, 39, including four newly annotated, localized to the proposed YDR (106–116 Mb) (Figure 7c, Figure S2, Table S5). Similarly, reciprocal BLAST searches against the *S. oleracea* YY reference genome (Sp_YY_v1), which has a well-characterized SDR subdivided into YDR, PAR1, and PAR2 (She *et al*., 2023), mapped 25 genes to the YDR. In *S. oleracea*, five previously unannotated gene models were predicted, three of which were located in the YDR (Figure S3, Table S5). PCR validation of eight randomly selected genes from YDR and PAR in male and female *S. turkestanica* genomes confirmed that YDR genes amplified exclusively in males, whereas PAR genes were detected in both sexes, supporting their predicted SDR locations (Figure S4, Table S7).

Transcriptomic profiling identified candidate genes for sex differentiation in *S. turkestanica*

### Differential expression of genes from *de novo* assembled SDR contigs

To identify potential regulators of sex differentiation within the SDR, we analyzed RNA-seq data from male and female *S. turkestanica* flowers across five developmental stages (Figure S5). Genes differentially expressed in at least two consecutive stages were considered valid DEGs. Pre-processed high-quality reads (average 13.9 million paired-end reads per sample; n = 30; Table S8) were aligned to the *de novo* assembled male-specific contigs. Stage-specific analysis identified 23 DEGs, seven of which mapped to the proposed YDR, including two newly annotated genes, *TU_SDR00082* and *TU_SDR000103* (Figure 8). Most of the sex-and stage-specific DEGs were nuclear and contained MYB, bZIP, or RNA-binding splicing factor domains (Table S9), known to regulate flower development and sex differentiation (Murase *et al*., 2017; Guo *et al*., 2025; Romera-Branchat *et al*., 2025). Given that sex is determined at early floral stages (Ma *et al*., 2022), we focused on genes exhibiting early-stage male-specific expression. Based on this analysis, *TU_SDR00075*, *TU_SDR00081*, *TU_SDR00082*, *TU_SDR00087*, *TU_SDR000174*, and *TU_SDR000177* were predicted as candidate sex-determining genes, while *TU_SDR000103* and *TU_SDR000168*, showing male-specific expression, may also contribute to sex differentiation (Figure 8, Table S9). Interestingly, *TU_SDR000115* and *TU_SDR000178* displayed higher expression in female flowers, whereas *TU_SDR00037*, *TU_SDR000108*, and *TU_SDR000148*, expressed at late male stages, may be involved in pollen development, maturation, and viability.

**Figure 8.**
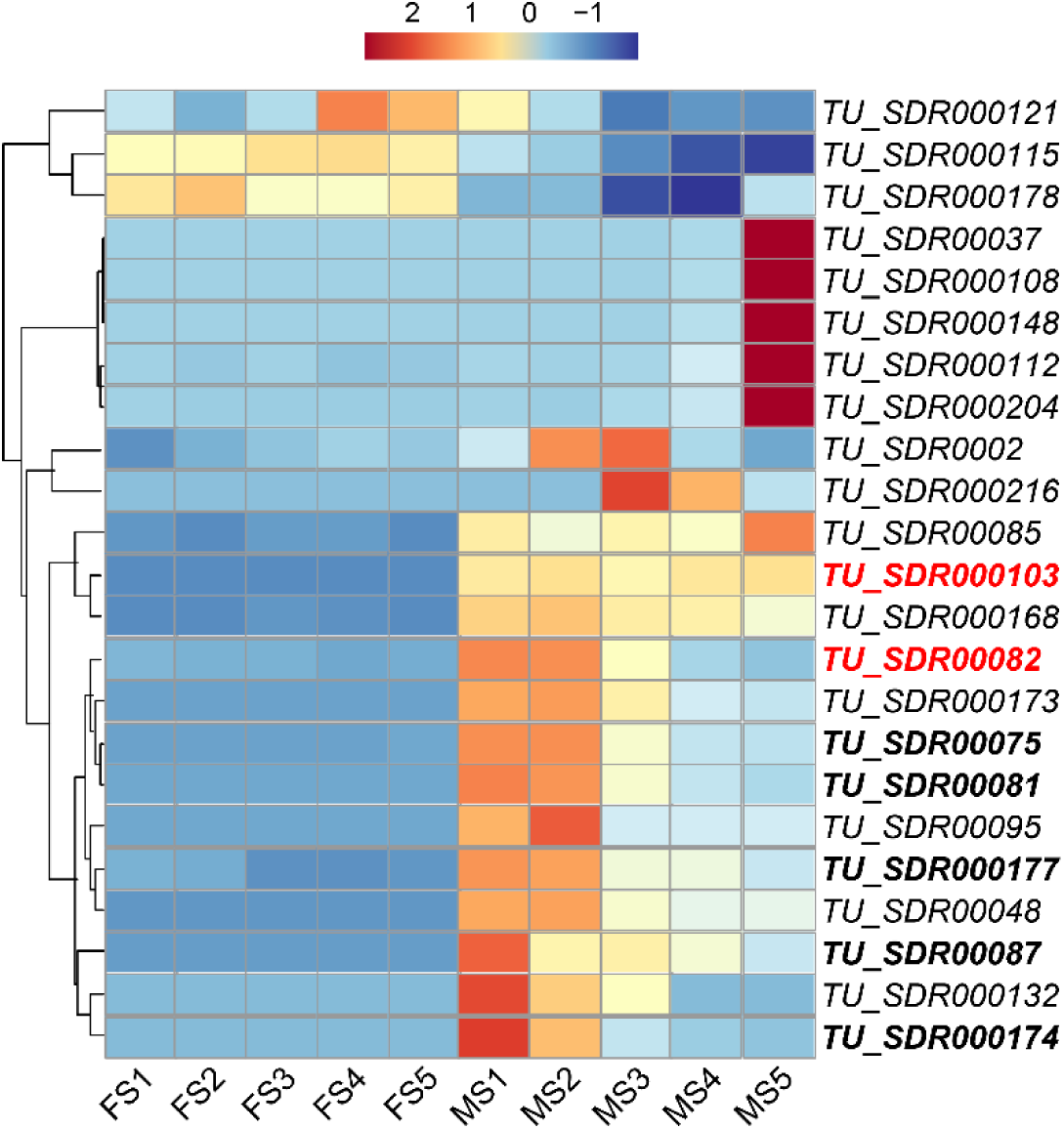
Temporal expression of *de novo* assembled SDR gene models in reproductive tissues. Heatmap of TPM based expression for sex-determining region (SDR) genes that are differentially expressed between male and female flowers across developmental stages. Genes are arranged by hierarchical clustering to reveal the temporal activation patterns. FS1–FS5 and MS1–MS5 denote female and male flower stages 1–5, respectively. Heatmap colors correspond to per-gene z-score. Gene IDs in bold are from the proposed Y-duplication region. newly predicted gene models are in red color.

### Differential expression of genes based on the reference genome

To gain a global perspective of gene expression during flower development, we mapped the same RNA-seq dataset to the *S. turkestanica* Tu17S31XY genome (She *et al*., 2025), incorporating nine newly annotated genes (Table S5). Principal component analysis (PCA) using genes with TPM >1 clearly separated male and female flowers, with the first component explaining 68.7% of variance (Figure 9a). Early-stage samples of both sexes clustered closer together, suggesting less pronounced transcriptomic differences at these stages. Overall, we identified 4,587 DEGs between male and female flowers, of which 3,735 were male-biased and 852 female-biased (Table S10). A stepwise increase in DEGs from stage 1 (n=130) to stage 5 (n=3,938) reflected the PCA trends (Figure 9b). Male-biased genes consistently outnumbered female-biased genes at each stage, and this pattern persisted when DEGs were categorized into autosomes versus the XY chromosomes.

**Figure 9.**
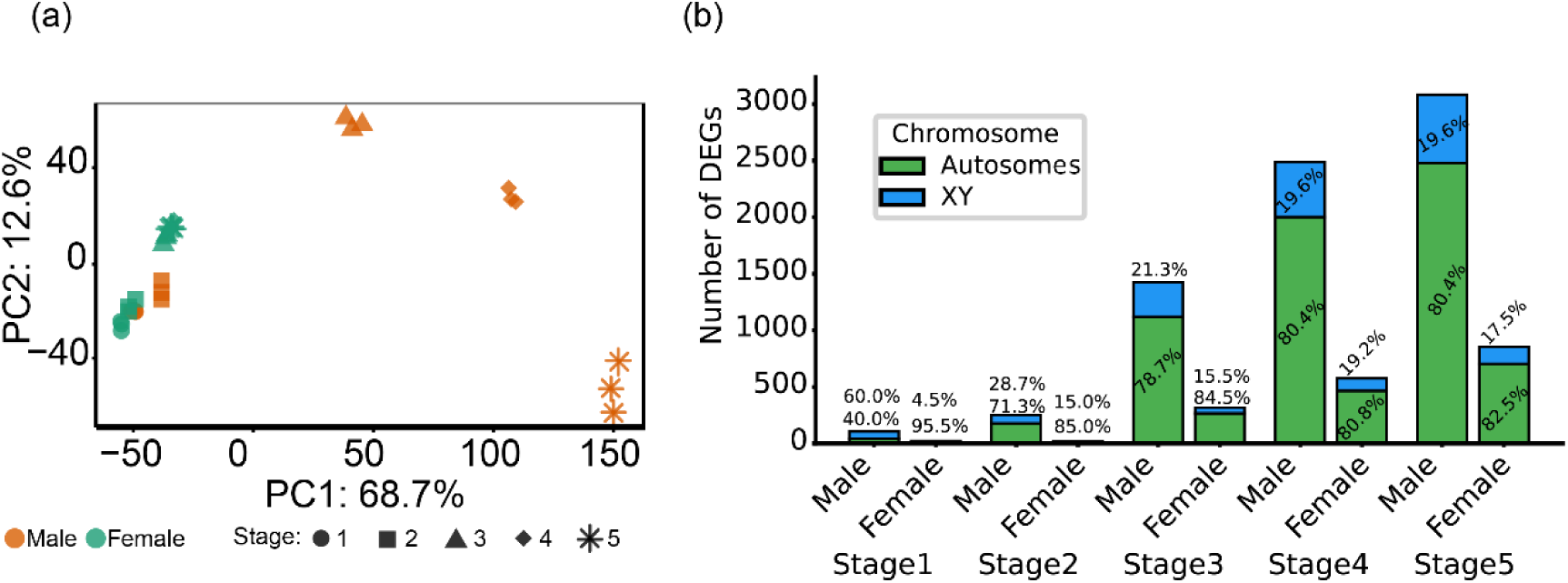
Gene expression dynamics of *S. turkestanica* male and female flowers. (a) principal component analysis plots of RNA-Seq data based on TPM. (b) The absolute number of DEGs at stage 1-5 of male and female flower development. The stacked bars also show the proportion of male and female biased genes on autosomes and XY chromosome.

GO enrichment analysis highlighted that male-biased genes, particularly in the SDR, were associated with metabolic processes, cell differentiation, reproduction, pollination, and flower development (Table S11), whereas female-biased genes were enriched for anatomical and post-embryonic development, reproduction, and flower development (Table S12). These results suggest a potential role of SDR-specific genes in sex determination and differentiation.

For downstream analyses, we focused on DEGs consistent across at least two consecutive developmental stages, reducing the dataset to 2,660 genes. Within the Tu17S31XY SDR, 34 DEGs were identified (Figure S6), 19 of which overlapped with DEGs from the *de novo* assembled SDR contigs and exhibited consistent expression patterns at both contig and genome levels. These findings strengthen the candidature of the target genes for sex determination.

### WGCNA identified SDR enriched modules associated with early-stage male flower development

To identify genes co-expressed with the candidate sex-determining genes, we performed Weighted Gene Co-expression Network Analysis (WGCNA) using the top 9,842 genes, clustering them into 15 modules (corr <0.8) based on expression patterns (Figure 10ab, Figure S7ab, Figure S8). We hypothesized that candidate sex-determining genes would be early-stage DEGs from the SDR region and members of modules associated with early male flower development. Among the 15 modules, ‘papayawhip’ (n=62), containing 40 DEGs including 21 from the SDR, was significantly correlated with early male stages (stages 1–2) (Figure 10ab, Figure S9, Table S13). GO enrichment was underpowered due to the small module size; therefore, we manually annotated ‘papayawhip’ members using InterProScan, UniProt, and TAIR, identifying transcription factors, transporters, metabolic enzymes, cell wall–related proteins, and RNA-binding proteins (Table S13). Another module, ‘bisque’ (n=1063), with 747 DEGs including four from the SDR, was significantly correlated with late male stages (Figure 10a, Figure S9, Table S14) and enriched for pollination, cell growth, and transport processes (Table S15). Although the minimum module size was set to 30, we retained the unique small module ‘aquamarine’ (n=17), which included five DEGs (one from the SDR) and was significantly correlated with early female stages (Figure 10a, Figure S9). Heat-shock proteins present in the ‘aquamarine’ module (Table S16).

**Figure 10.**
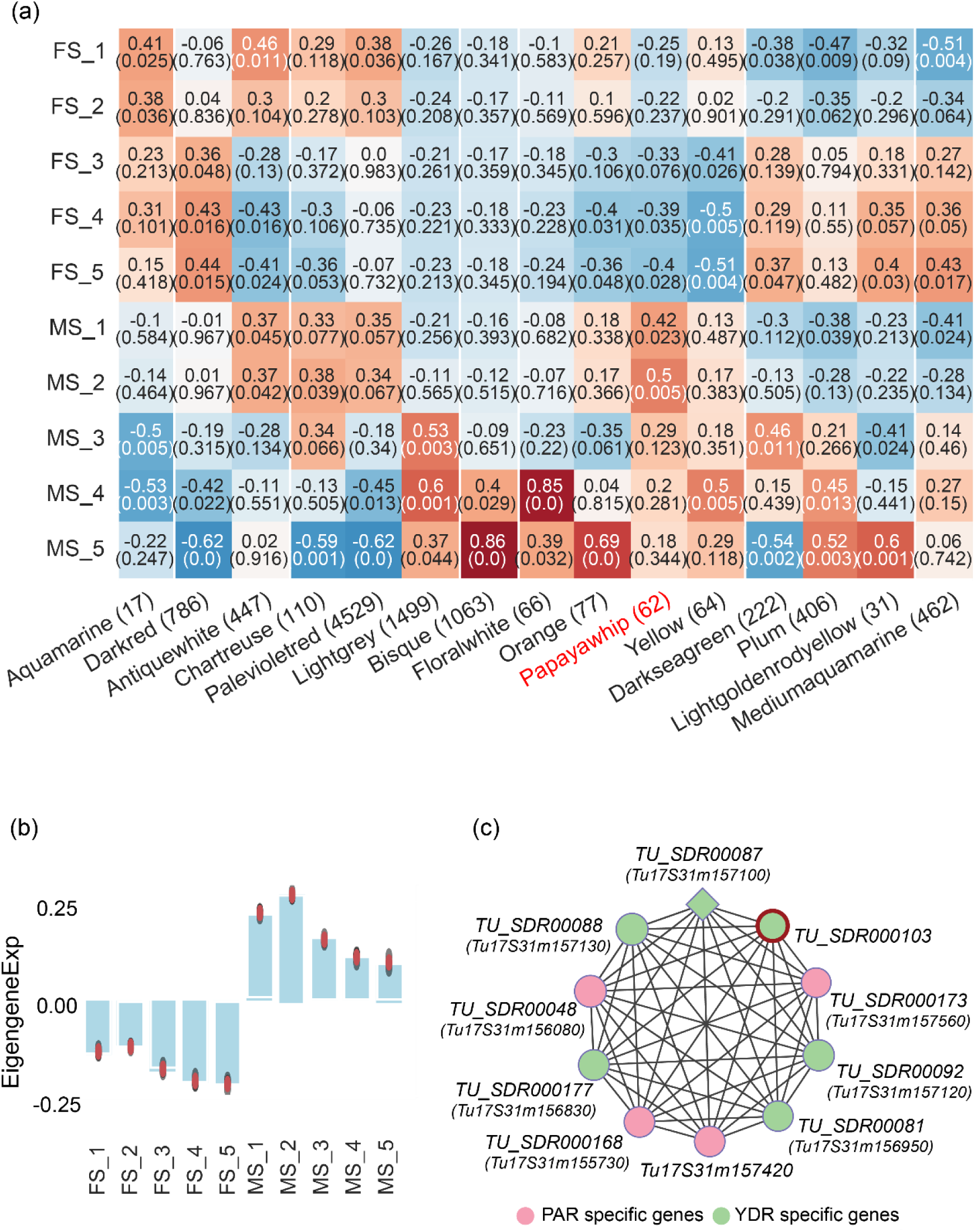
Weighted co-expression network analysis of *S. turkestanica*. (a) Heatmap showing correlations between gene co-expression modules and male (MS1–MS5) and female (FS1–FS5) flower developmental stages. The X-axis indicates module names and their corresponding gene counts, while the Y-axis represents developmental stages. Target module ‘papyawhip’ is in red. (b) Bar plot of eigengene expression for the target module ‘papayawhip,’ which is associated with early male flower development. (c) Network visualization of hub gene (diamonds) within the ‘papayawhip’ module. Newly identified genes in this study are highlighted with red circle. Gene IDs in brackets are from the *S. turkestanica* published genome. Node size reflects intramodular connectivity (kWithin), and edge thickness represents connection strength (weight) between genes.

### Network graph visualization

As the module of primary interest, intramodular connectivity analysis of the ‘papayawhip’ identified the YDR-specific, bZIP domain containing, candidate sex-determining gene *TU_SDR00087* as a potential hub (Figure 10c, Table S13). Notably, five additional candidate sex-determining genes (*TU_SDR00081, TU_SDR00087, TU_SDR000103, TU_SDR000168,* and *TU_SDR000177*) ranked among the top nine genes most strongly connected to the hub, indicating that SDR-specific DEGs are tightly co-expressed within ‘papayawhip’ module.

### qPCR validation of candidate sex-determining gene expression

To validate our RNA-seq and WGCNA findings, selected candidate DEGs were analyzed by qPCR across five floral developmental stages in male and female flowers. Genes were chosen based on their localization in the SDR, male-specific RNA-seq expression patterns, predicted nuclear localization, protein domains linked to flower development, and membership in the target co-expression module (Figure 8, Figure S6, Table S9, Table S13). All YDR-localized genes exhibited Ct values around 35 in female flower samples, indicating no detectable expression. *TU_SDR00081*, *TU_SDR00087*, *TU_SDR000168*, and *TU_SDR000174* showed significantly higher expression at early stages (Stages 1–2) of male flower development, whereas the newly annotated *TU_SDR00082* and *TU_SDR000103* exhibited consistent male-specific expression (Figure 11b-f, i-j). In contrast, the expression level of *TU_SDR000037*, *TU_SDR000108*, and *TU_SDR000148* was significantly higher at the late stage (stage 5) of male flower development (Figure 11agh).

**Figure 11.**
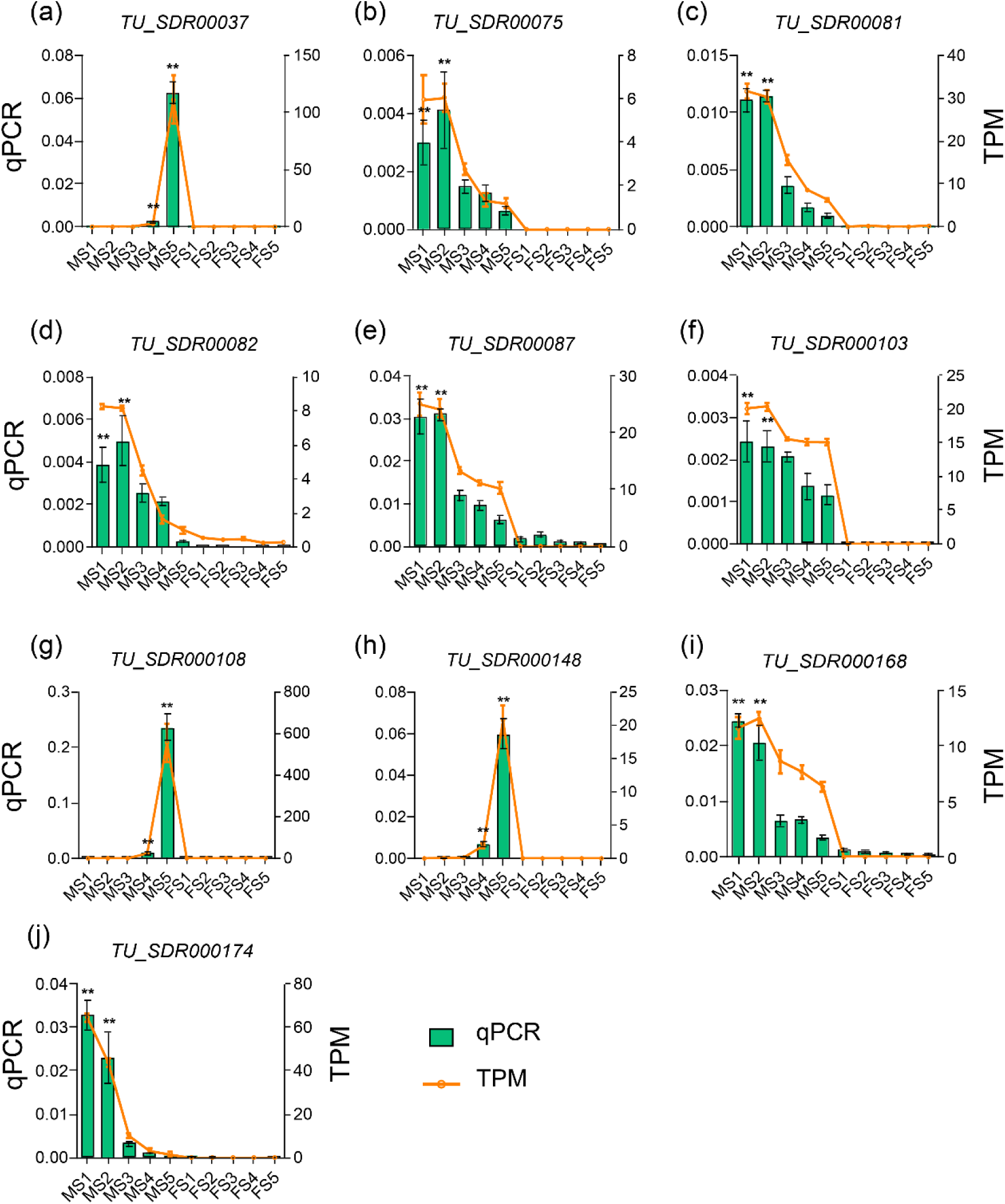
Expression analysis of selected SDR genes across five developmental stages in *S. turkestanica* male and female flowers. *spUBI* served as the internal reference gene. Two technical and three biological replicates were used to reduce error. Error bars represent standard deviation (n = 3). Statistical significance was assessed by Student’s t-test comparing early male stages (MS1–MS2) versus all other stages (MS3–MS5 and FS1–FS5): *p < 0.05; **p < 0.01. MS1–MS5 and FS1–FS5 denote male and female flower developmental stages 1–5, respectively. qPCR results are shown as bar graphs, while RNA-seq (TPM) values of the same genes are plotted as line graphs on the secondary Y-axis to validate expression patterns.

### Candidate genes for sex determination

Sex differentiation in plants is initiated early during floral development (Ma *et al*., 2022); therefore, we hypothesize that candidate sex-determining genes should be single-copy, located within the sex-determining region (SDR), and exhibit male-specific expression during early floral stages. According to these criteria, we propose *TU_SDR00087* and *TU_SDR000174*, both located in the Y duplication region (YDR), as strong candidates for sex determination in *S. turkestanica*. Additionally, *TU_SDR000168*, located in the pseudo-autosomal region (PAR1), may function as a potential modulator of sex differentiation (Figure 8, Figure 10c, Figure 11eij, Figure S6, Table S9).

## Discussion

Sexual dimorphism, characterized by the existence of distinct male and female individuals, is common in animals but occurs in only about 6% of terrestrial plant species (You *et al*., 2025). Understanding the genetic basis of dioecy is essential, particularly in economically important crops that exhibit this trait (Shi *et al*., 2025). Within the *Spinacia* genus, *S. turkestanica* is the closest wild relative of *S. oleracea*, a dioecious species widely cultivated as a leafy vegetable in China (She *et al*., 2025). This evolutionary relationship makes *S. turkestanica* a valuable model for investigating the genetic mechanisms underlying sex determination. Notably, the master regulator(s) of sex determination in *Spinacia* remain unidentified (Li *et al*., 2022). To address this knowledge gap, we integrated k-mer–based *de novo* SDR assembly, transcriptomic profiling, and weighted gene co-expression analysis (WGCNA) across five developmental stages of male and female flowers, enabling the identification of candidate sex-determining genes in *S. turkestanica*.

### Spinach is XY system with SDR categorized into YDR and PAR

Several studies indicate that the SDR typically comprises male-specific duplicated regions on the Y chromosome, flanked by recombination-suppressed boundaries (Liao *et al*., 2020; She *et al*., 2024; Shi *et al*., 2025). Although the SDR in the published *S. turkestanica* male genome has not yet been explicitly divided into YDR and PAR (She *et al*., 2025), k-mer–based mapping of sex-specific PE reads (Figure 7, Figure S1a), complemented by reciprocal BLASTp analysis of *de novo* assembled SDR contigs against the *S. oleracea* (Sp_YY_v1) genome (Table S5) and PCR validation (Figure S4), suggests the presence of a similar structural organization in *S. turkestanica*. Furthermore, *S. turkestanica* exhibits XY system of sexual dimorphism, wherein sex-associated SNPs are consistently heterozygous in males and homozygous in females (Figure S1b, Table S2), mirroring the SNP heterozygosity patterns reported for *S. oleracea* (She *et al*., 2023).

### K-mer analysis reveals novel SDR associated sequences

To identify the sex-determining region (SDR) in *S. turkestanica*, we employed a reference-independent k-mer–based approach. While the SDR in *S. turkestanica* is located on chromosome 4 (She *et al*., 2025), our k-mer analysis revealed additional SDR-associated sequences residing in previously unassembled or unanchored contigs of the published *S. turkestanica* genome (Figure 7b,d). This methodology, validated in other dioecious species like kiwifruit (Akagi *et al*., 2019), *Ginkgo biloba* (Liao *et al*., 2020) grapevine (Massonnet *et al*., 2020), *Morus* plants (Xia *et al*., 2022), and hemp (Shi *et al*., 2025), proved particularly valuable for *S. turkestanica*, where traditional synteny-based approaches using the *S. oleracea* Sp_YY_v1 genome (She *et al*., 2023) had failed to capture these unanchored genomic elements. The reference-free nature of the k-mer strategy enabled a more comprehensive characterization of the SDR, effectively overcoming limitations posed by assembly gaps in existing genomic resources (Roberts *et al*., 2025).

### Repeat-rich architecture of the assembled SDR

We successfully assembled a ∼21.5 Mb sex-determining region (SDR) in *S. turkestanica*, with approximately 84% of the sequence composed of repetitive elements (Table S4). SDRs are known to evolve through the duplication of autosomal regions, followed by sequence divergence, leading to the accumulation of repeats and a reduction in gene density (She *et al*., 2023, 2024). In line with this evolutionary pattern, the SDRs in the published genomes of *S. turkestanica* and *S. oleracea* harbor 85.05% and 86.04% repetitive content, respectively (She *et al*., 2025). Our findings are consistent with these reports and further underscore the highly repetitive nature of the spinach SDR.

Gene modeling and co-expression analysis reveal candidate sex-determining genes Genes located within the SDR and showing differential expression between male and female plants during early floral development are likely to play crucial roles in sex differentiation and determination (Ma et al., 2022). Using the MAKER annotation pipeline, we predicted a total of 226 gene models within the *S. turkestanica* SDR, including nine newly identified high-confidence genes (Table S5).

Comparative transcriptomic and co-expression (WGCNA) analyses of male and female floral tissues across developmental stages have been effective in pinpointing candidate sex-determining genes in several dioecious species (You *et al*., 2025). For instance, in papaya, RNA-seq analyses of early floral stages uncovered several differentially expressed transcription factors involved in sex differentiation (Liu *et al*., 2021). Similarly, in hemp, transcriptome profiling coupled with WGCNA of vegetative and reproductive tissues from both sexes identified multiple candidates (Shi *et al*., 2025).

Our integration of RNA-seq, WGCNA, and qPCR validation uncovered several promising candidate sex-determining gene in *S. turkestanica*. Notably, the RNA-binding splicing factor domain-containing gene *TU_SDR000168* from the proposed PAR, the bZIP domain-containing *TU_SDR000087*, and the MYB domain-containing *TU_SDR000174* from the proposed YDR, were all significantly upregulated and co-expressed during early stages (stage 1–2) of male flower development (Figure 8, Figure 10c, Figure 11, Figure S6). These genes represent strong candidates for further functional characterization in the context of sex determination.

Serine/arginine-rich splicing factor may regulate the candidate sex determining gene/s Alternative splicing enables the generation of multiple mRNA isoforms from a single gene, expanding transcriptomic and proteomic diversity (Park *et al*., 2018). This process is tightly regulated by distinct classes of RNA-binding proteins, including serine/arginine-rich (SR) proteins, heterogeneous nuclear ribonucleoproteins (hnRNPs), and tissue-specific splicing factors (Guo *et al*., 2025). Among the identified candidates, *TU_SDR000168*, an SR domain-containing gene located in the proposed PAR (Table S5) of *S. turkestanica*, was significantly upregulated during early stages of male floral development (Figure 8, Figure 11i).

In *Arabidopsis thaliana*, the SR protein SR45 modulates the alternative splicing of genes involved in floral transition. Loss of *SR45* function leads to upregulation of *FLOWERING LOCUS C* (*FLC*) and delayed flowering (Albaqami *et al*., 2019). Similarly, the *CDKG1* gene, another SR-type splicing factor, is essential for pollen development; its loss results in defective pollen wall formation and male sterility (Nibau *et al*., 2020). Furthermore, isoforms of *AUXIN RESPONSE FACTOR 8* (*ARF8*), particularly *ARF8.1*, have been shown to directly regulate key transcription factors such as *TDF1*, *AMS*, and *MS188*, underscoring the importance of alternative splicing in male reproductive development (Ghelli *et al*., 2023).

Functional annotation using InterProScan database classified *TU_SDR000168* as a member of the “transformer-2 sex-determining protein-related” superfamily. The *tra-2* gene, characterized by RS-rich motifs, is well-known in insects for its pivotal role in sex determination through regulation of sex-specific splicing (Mattox *et al*., 1990; Laohakieat *et al*., 2020). Based on these findings, we hypothesize that *TU_SDR000168* may produce male-specific isoforms of key downstream genes involved in the sex differentiation pathway in spinach. However, its precise functional role remains to be validated through further molecular and genetic analyses in *S. turkestanica*.

### Y-linked bZIP transcription factor as a putative sex-determining gene

Basic leucine zipper (bZIP) transcription factors are widely reported in plants and are known to regulate diverse processes including growth, development, morphogenesis, and signal transduction (Han *et al*., 2023*a*). In this study, we identified a YDR-specific bZIP gene, *TU_SDR00087*, showing male-specific expression and hub status in the ‘papayawhip’ module, which is strongly associated with early male flower development in *S. turkestanica* (Figure 8, Figure 10c, Figure 11e). Orthologs of this gene were also detected in the YDR of *S. oleracea* and *S. tetrandra* (She *et al*., 2025). Consistent with our findings, the ortholog *YY_141140.1* in *S. oleracea* is also expressed exclusively in male flowers at early developmental stages (She *et al*., 2023).

In *Arabidopsis*, the ortholog of *TU_SDR00087* is known as *FD* (AT4G35900), a bZIP transcription factor that functions as a core component of the florigen activation complex (FAC), which promotes flowering and regulates floral organ identity (Abe *et al*., 2005; Romera-Branchat *et al*., 2025). Similarly, in rice, *OsFD4*, another member of the bZIP family, acts within the FAC to facilitate flowering at the shoot apical meristem (Cerise *et al*., 2021). Taken together, these observations suggest that *TU_SDR00087* may play a pivotal role in male flower development and potentially act as a key regulator in the sex-determination pathway of spinach.

### MYB genes may regulate distinct stages of male flower development

MYB transcription factors (TFs) are known to play pivotal roles in male reproductive development, including tapetum formation, pollen maturation, filament elongation, and regulation of downstream genes involved in floral organogenesis (Wang *et al*., 2023; Zhang *et al*., 2024). In our study, two MYB-domain–containing genes, *TU_SDR000174* and *TU_SDR000108*, exhibited distinct expression patterns during male flower development in *S. turkestanica*.

YDR specific *TU_SDR000174* encodes a RADIALIS-like (RL) MYB-related TF that showed significantly higher expression during the early stages (stage 1-2) of male floral development (Figure 8, Figure 11j). While RAD-like genes are associated with establishing floral asymmetry (Baxter *et al*., 2007), recent studies have uncovered additional functions. For example, ectopic overexpression of *DkRAD*, a RAD-like gene from hexaploid persimmon, in Arabidopsis and tobacco induced excessive gynoecium growth and was termed a “restorer of femaleness” (Masuda *et al*., 2022). These findings suggest that *TU_SDR000174* may have similarly acquired a novel role in sex differentiation in spinach.

Interestingly, in *Asparagus officinalis*, a male genome-specific MYB gene, *MSE1* (also referred to as *AoMYB35*), exhibits elevated expression during early flower development (Murase *et al*., 2017). Knockout of its Arabidopsis ortholog leads to male sterility. *AoMYB35* expression is restricted to the stamen at the premeiotic tapetum development stage, further supporting a role for MYB genes in sex-specific processes (Tsugama *et al*., 2017).

In contrast, PAR localized, *TU_SDR000108* showed peak expression at the late stage (Stage 5) of male flower development (Figure 8, Figure 11g). Its ortholog *YY_141510.1* in *S. oleracea* showed a similar temporal expression pattern (Zhang *et al*., 2024). Known Arabidopsis homologs include *MYB101*, involved in pollen tube reception and expressed in mature pollen grains and tubes (Liang *et al*., 2013), and *MYB65*, which regulates programmed cell death in the tapetum during anther development (Millar and Gubler, 2005). These parallels suggest that *TU_SDR000108* may be critical for pollen viability, maturation, and fertilization in spinach.

Collectively, these findings support stage-specific roles for MYB transcription factors in male floral development and reinforce their potential contribution to sex differentiation in *S. turkestanica*.

### SDR Gene expressing at later stages of flower development may regulate the pollen development

A cupin domain-containing gene, *TU_SDR000148*, was significantly expressed during the late stages of male flower development (Figure 8, Figure 11h). Its Arabidopsis homolog, *AT2G28680*, is specifically expressed in stamen tissue. Similarly, *GhGLP4*, a cupin domain gene in *Gossypium hirsutum*, has been shown to play a crucial role in stamen development. RNAi-mediated silencing of *GhGLP4* resulted in reduced anther number and shortened stamens (Zheng *et al*., 2021), highlighting the potential involvement of cupin domain genes in pollen development.

Another late-stage expressed gene, *TU_SDR00037*, contains a Domain of Unknown Function (DUF868) and showed male-specific expression during the late stages of flower development (Figure 8, Figure 11a). This gene is not annotated in the *S. oleracea* Sp_YY_v1 genome yet (Table S9). In the Pfam 35.0 database, 24% of the gene families are tagged as DUF type and their function yet to be explored (Lv *et al*., 2023). While DUF868 has not yet been functionally characterized, *TU_SDR00037* localizes to the nucleus, and its *Arabidopsis* homolog, *AT2G04220*, is known to be expressed in mature pollen tissue. Together, these findings suggest a possible role for *TU_SDR000148* and *TU_SDR00037* in pollen development and maturation.

## Conclusion

In this study, we identified 23 differentially expressed genes (DEGs) between male and female floral tissues of *S. turkestanica*. Based on SDR localization, conserved domain, early stage-specific co-expression during flower development, we propose *TU_SDR00087*, *TU_SDR000168*, and *TU_SDR000174* as strong candidate sex-determining genes. While these genes are likely contributors to sexual dimorphism in *S. turkestanica*, their roles as bona fide sex-determination factors remain to be experimentally validated. Future functional studies, including gene knockout and overexpression analyses, will be critical to elucidate their precise involvement in the sex determination pathway

## Supplementary data

Figure S1: Identification of *S. turkestanica* YDR and validation of XY system.

Figure S1. Integrated transcriptomic view of the newly identified gene models in *S. turkestanica*.

Figure S3. Integrated transcriptomic view of the newly identified gene models in *S. oleracea*.

Figure S2: PCR based validation of *S. turkestanica* YDR.

Figure S3. Five stages of *S. turkestanica* male and female flowers development.

Figure S6. Comparative heatmaps of candidate SDR genes at contig and reference genome level.

Figure S7. Quality control analysis of the sample data and the WGCNA modules. Figure S4. Hierarchical clustering of *S. turkestanica* Tu17S31XY genes.

Figure S5.WGCNA based module eigengene heatmap.

Table S1: Male and female specific PE reads depth in *S. turkestanica* published genome. Table S2. Results of GWAS for sex in *S. turkestanica*.

Table S3: Contigs assembly statistics. Table S4: Repetitive sequences statistics.

Table S5. Gene annotation of the *de novo* assembled sex determining region of *S. turkestanica*.

Table S6: Gene model statistics.

Table S7. List of PCR primers used to validate the Y-duplication region in *S. turkestanica*.

Table S8. Quality assessment statistics of transcriptome data.

Table S9. Functional annotation of DEGs from the *de novo* assembled SDR contigs of *S. turkestanica*.

Table S10. Mean TPM values per flower development stage of *S. turkestanica*.

Table S11. GO enrichment analysis of *S. turkestanica* up regulated DEGs.

Table S12. GO enrichment analysis of *S. turkestanica* down regulated DEGs.

Table S13. Genes members of male-specific early stage correlated modules ’papayawhip’ in weighted gene co-expression network analysis (WGCNA).

Table S14. Genes members of male-specific late stage correlated modules ’bsique’ in weighted gene co-expression network analysis (WGCNA).

Table S15. GO enrichment analysis of male-specific late stage correlated modules ’bsique’.

Table S16. Genes members of female-specific early stage correlated modules ’aquamarine’ in weighted gene co-expression network analysis (WGCNA).

Table S17. List of qPCR primers used in this study.

## Author contributions

SM: writing - original draft, conceptualization; data curation; SL, TZ, XL, LY: formal analysis; AA-M: review & editing; NL: conceptualization, review & editing; CD: conceptualization, funding acquisition, supervision.

## Conflict of interest

No conflict of interest declared.

## Funding

This work was supported by the Natural Science Foundation of Henan Province (General Program, Grant No. 242300420162 and 250300420203), the Henan Province International Science and Technology Cooperation Project (Grant No. 252102520005), the Henan Province University’s Engineering and Technology Center of Conservation and Utilization for Genuine Chinese Medicinal Herbs, and Xinxiang Engineering and Technology Center of Conservation and Utilization for Chinese Medicinal Herbs.

## Data availability

The raw sequence data reported in this paper have been deposited in the Genome Sequence Archive in National Genomics Data Center, China National Center for Bioinformation / Beijing Institute of Genomics, Chinese Academy of Sciences (GSA: CRA031114) that are publicly accessible at https://ngdc.cncb.ac.cn/gsa

## Notes

### Competing Interest Statement

The authors have declared no competing interest.

